# Feedback in the β-catenin destruction complex imparts bistability and cellular memory

**DOI:** 10.1101/2022.01.28.478206

**Authors:** Mary Jo Cantoria, Elaheh Alizadeh, Janani Ravi, Nawat Bunnag, Arminja N. Kettenbach, Yashi Ahmed, Andrew L Paek, John J. Tyson, Konstantin Doubrovinski, Ethan Lee, Curtis A. Thorne

## Abstract

Wnt ligands are considered classical morphogens, for which the strength of the cellular response is proportional to the concentration of the ligand. Herein, we show an emergent property of bistability arising from feedback among the Wnt destruction complex proteins that target the key transcriptional co-activator β-catenin for degradation. Using biochemical reconstitution, we identified positive feedback between the scaffold protein Axin and the kinase GSK3. Theoretical modeling of this feedback between Axin and GSK3 predicted that the activity of the destruction complex exhibits bistable behavior. We experimentally confirmed these predictions by demonstrating that cellular cytoplasmic β-catenin concentrations exhibit an “all-or-none” response with sustained memory (hysteresis) of the signaling input. This bistable behavior was transformed into a graded response and memory was lost through inhibition of GSK3. These findings provide a mechanism for establishing decisive, switch-like cellular response and memory upon Wnt pathway stimulation.

**One Sentence Summary:** Positive feedback within the β-catenin destruction complex gives rise to bistability and memory in response to Wnt stimulation, imparting signal transduction accuracy and insulation.

## RESULTS

Wnt/β-catenin signaling is involved in organism development, stem cell maintenance and is misregulated in human disease. At the core of this signaling pathway is the β-catenin destruction complex, comprised of the kinases glycogen synthase kinase 3 (GSK3) and casein kinase 1 alpha (CK1α), and the scaffolding proteins Axin and Adenomatous polyposis coli (APC). In the absence of Wnt ligands, phosphorylation of the transcriptional co-activator β-catenin within the destruction complex targets β-catenin for ubiquitin-mediated proteasomal degradation, thereby maintaining low levels of cytoplasmic and nuclear β-catenin. Wnt signaling inhibits phosphorylation of β-catenin to block its turnover; accumulated β-catenin subsequently enters the nucleus to mediate a Wnt-specific transcriptional program required for animal development and tissue homeostasis (*1*).

Although Wnt ligands are considered classical morphogens, Wnt gradients are dispensable for proper patterning during development in some contexts (*2-4*). To better understand the biochemical function of the β-catenin destruction complex and to assess how critical steps within the complex impact behavior of the Wnt pathway, we performed biochemical reconstitutions of the destruction complex with Xenopus egg extracts and purified proteins. Based on these measurements, we developed mathematical simulations of destruction complex dynamics and validated our model by performing single-cell analyses of β-catenin behavior.

Previous studies in cultured mammalian cells and *in vitro* reconstitution have shown that the scaffold protein Axin is a direct target of GSK3 (*5, 6*). Because *Xenopus* egg extracts are readily amenable to biochemical studies and faithfully recapitulate signaling dynamics that control β-catenin turnover (*7*), we examined the regulation of Axin by GSK3 in extracts (Fig. 1A). Consistent with previous studies (*5*), inhibition of GSK3 with LiCl induced Axin turnover (Fig. 1B,C). Stabilization of Axin required both the GSK3 phosphorylation sites at serine 322 and serine 326, and the GSK3 binding site (GBS) on Axin (Fig. 1C) (*8*).

**Fig. 1.**
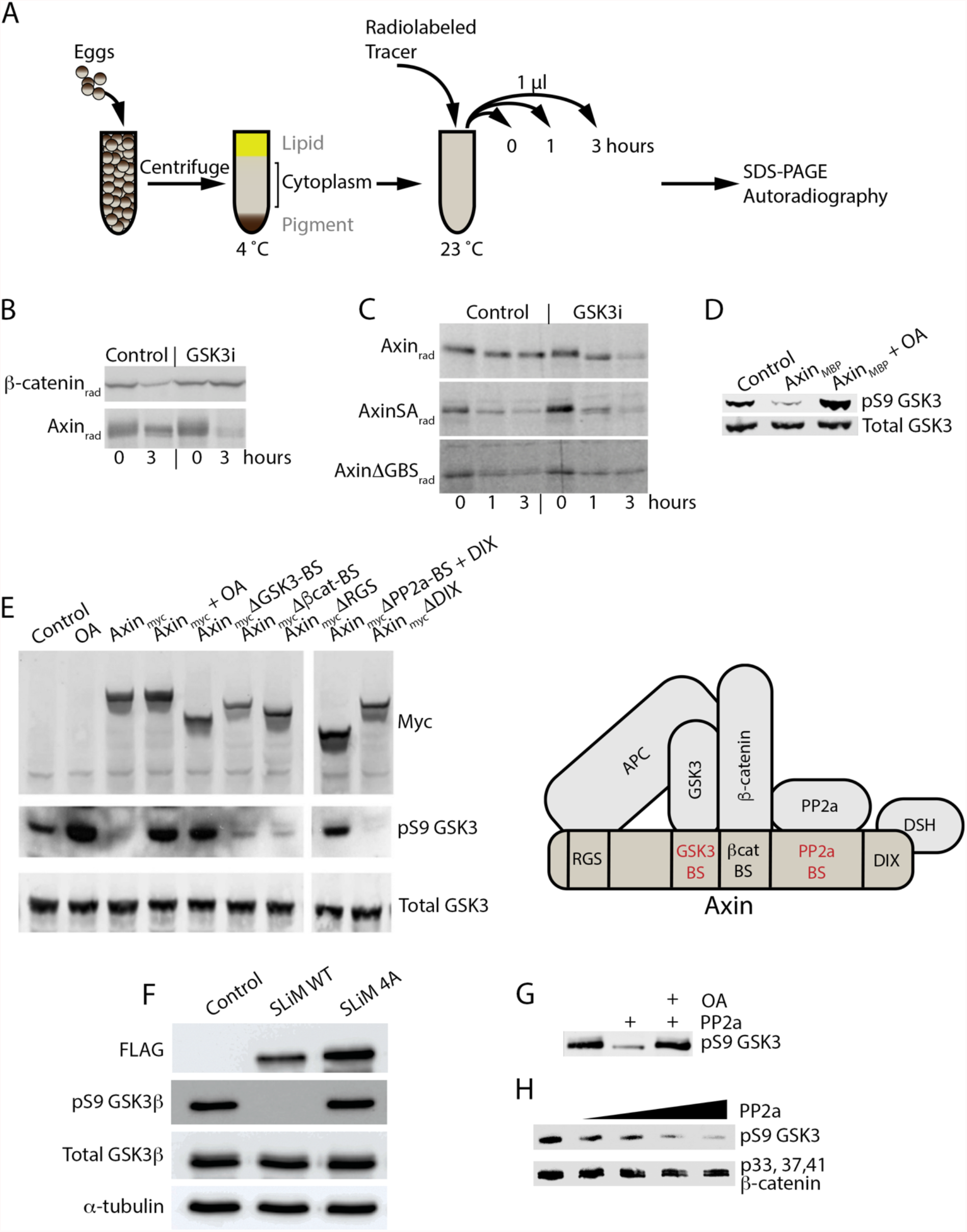
GSK3 and Axin mutually activate in Xenopus extracts and mammalian cells. (**A**) Experimental scheme. Cytoplasmic fraction of *Xenopus* egg extracts. Extracts are collected, spiked with radiolabeled (rad) [^35^S] β-catenin or [^35^S] Axin, and aliquots are removed at the indicated time points for analysis by SDS-PAGE and autoradiography. (**B**) Turnover of radiolabeled [^35^S] β-catenin or [^35^S]Axin in Xenopus extracts. LiCl (GSK3i) and NaCl (Control) (50 mM each) were added to extracts as indicated. (**C**) As in (B), turnover of Axin, AxinSA (serine 322 and 326 mutated to alanine), and AxinΔGBS (GSK3 binding site) in Xenopus extracts. (**D**) Axin promotes dephosphorylation of pS9 GSK3 in Xenopus extract, which is blocked by OA (200 nM). MBP-Axin (10 nM) was added to egg extract in the presence or absence of OA (10 nM), and pS9 GSK3 and GSK3 were detected by immunoblotting. (**E**) The β-catenin binding site, APC binding site, and DIX domain are dispensable for Axin-mediated dephosphorylation of pS9 GSK3. Myc-tagged Axin truncation mutants were transfected into HEK293 cells, as indicated, and immunoblotting was performed. For OA treatment, cells were incubated with 10 nM OA for 2 hrs prior to lysis. (**F**) Expression of FLAG-tagged wild-type Axin (Axin SLiM WT) and FLAG-tagged Axin with mutations in the conserved B56 binding site that prevent the interaction of B56 with Axin (Axin SLiM 4A) in HEK293. (**G**) Reconstitution of pS9 GSK3 dephosphorylation by PP2A in the presence of Axin. Recombinant Axin (1 μM) and GSK3 (10 μM) were incubated with ATP for 30 min to allow for the autophosphorylation of GSK3. PP2A (1 μM) and OA (10 nM) were added for an additional 30 min, and samples were immunoblotted for pS9 GSK3. (**H**) PP2A preferentially dephosphorylates pS9 GSK3 versus β-catenin in the presence of Axin. The reaction was performed as in (F) but with the addition of β-catenin (10 μM).

As Axin is the limiting component of the destruction complex, overexpression of Axin promotes β-catenin degradation and inhibits Wnt signaling even in the absence of APC (*9, 10*). The limiting concentration of Axin provides a simple means for insulating a discrete pool of GSK3 that specifically targets β-catenin for phosphorylation (*10*). In addition, given its role as a scaffold, Axin is ideally positioned to regulate the activity of GSK3, thereby promoting both Axin stability and β-catenin degradation. We initially examined GSK3 activity in *Xenopus* egg extracts using a phospho-specific antibody that recognizes GSK3β phosphorylation at serine 9 (pS9 GSK3), which limits GSK3β activity. The addition of recombinant Axin to extracts resulted in a marked reduction in pS9 GSK3 (Fig. 1D). The requirement for a phosphatase in β-catenin degradation has been reported (*11*). Thus, we tested the effect of the phosphatase inhibitor okadaic acid (OA) on pS9 GSK3. We found that OA prevented the Axin-mediated reduction of pS9 GSK3 in *Xenopus* extracts (Fig. 1D), suggesting an OA-sensitive phosphatase requirement at this regulatory step.

To identify Axin regions that bind co-factors necessary for pS9 GSK3 dephosphorylation, we performed domain deletion analysis by expressing Axin mutants in HEK 293 cells (Fig. 1E). As expected, full-length Axin promoted loss of the inhibitory phosphorylation of GSK3, and OA blocked this effect, suggesting phosphatase dependence of GSK3 activation. Similarly, the deletion of the GSK3 binding site (GBS) or the phosphatase 2A (PP2A) domains of Axin prevented Axin-mediated inhibition of GSK3 phosphorylation, suggesting these regions are essential for GSK3 activation by Axin. In contrast, Axin lacking its β-catenin binding site (βcat-BS), APC binding site (RGS), or DIX domain still promoted the removal of the inhibitory serine 9 phosphorylation on GSK3; thus, these sites are not required for Axin-mediated removal of serine 9 phosphorylation on GSK3. Additionally, Axin was recently shown to contain a short linear motif (SLiM) that interacts with the B56 subunit of PP2A (*12, 13*). We made alanine mutants of this conserved SLiM sequence (Fig. S1) and found that SLiM 4A Axin mutants could not remove pS9 on GSK3 (Fig. 1F).

To test if PP2A could directly act on pS9 GSK3, we performed *in vitro* reconstitution using purified components of the destruction complex. We found that PP2A exhibited a preference for pS9 GSK3 (Fig. 1G) versus the β-catenin sites phosphorylated by GSK3 (phospho-serine 33, serine 37, and threonine 41; Fig. 1H). Based on these findings, we propose the following model: the majority of cytoplasmic GSK3 is in or fluctuating as the pS9 GSK3 state, which normally limits its activity. Upon pS9 GSK3 binding to Axin, pS9 GSK3 is targeted for dephosphorylation by Axin-bound PP2A. Dephosphorylated GSK3 is active and phosphorylates Axin to promote its stabilization. Active, dephosphorylated GSK3 and phosphorylated Axin (bound to APC) comprise a destruction complex state that is “fully activated” to phosphorylate β-catenin, targeting it for ubiquitin-mediated proteasomal degradation.

We built a theoretical model based on our biochemical observations to better understand the reaction kinetics within the β-catenin destruction complex (Fig. 2A). GSK3 concentration was kept constant as it is predicted to be degraded at a relatively slow rate (*10*). Rates of Axin synthesis and degradation were based on our *Xenopus* extract data and previous work (*10*). We translated our model (Fig. 2A) into a set of ordinary differential equations (ODEs) (TableS1) and solved them numerically and analytically in steady-state conditions (Fig. S2). The reaction rates and rate constants used in the model are listed in Table S2 and Table S3, respectively. As shown in Fig. 2A, our model showed a positive feedback loop between Axin and GSK3.

**Fig. 2.**
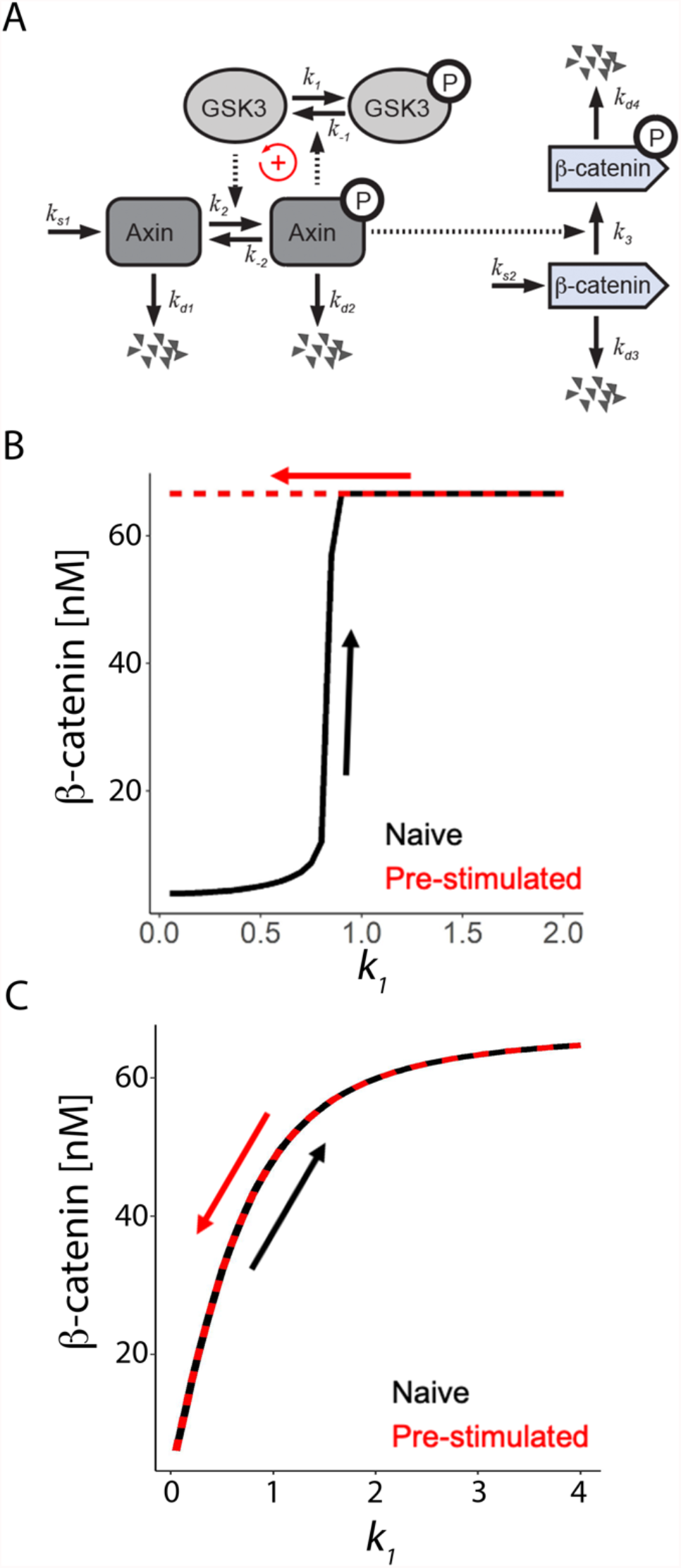
Mathematical modeling of the core β-catenin destruction complex components give rise to bistable Wnt activity. (**A**) Wiring diagram of β-catenin destruction complex feedback. Phosphorylated forms are denoted with “P.” The model consists of a positive feedback loop between GSK3 and Axin^p^. (**B**) We assume that an input Wnt signal changes the rate constant (*k*_*1*_) in the phosphorylation flux of GSK3 (*18*). The model shows bistable response in β-catenin. **(C)** Bistability is lost when GSK3 is dephosphorylated by a phosphatase activity that is independent of Axin^p^. Consequently, a graded β-catenin response is observed.

The function of the destruction complex is to promote the phosphorylation and subsequent ubiquitin-mediated degradation of β-catenin. When Axin^p^ and GSK3 are high, β-catenin is low, and the pathway is “off.” When Axin^P^ and GSK3 are low (e.g., via Wnt activation), cytoplasmic and nuclear β-catenin is high, and the pathway is “on.” Consistent with previous work, we modeled the Wnt signal to act on the active, destruction complex-bound GSK3 by directly increasing the inhibitory rate (k_1_) (*14*) and calculated the steady-state concentration of β-catenin. To model pathway activation, we started initially with a low value of k_1_, which was followed by a gradual increase. For each k_1_ value, the β-catenin concentration was determined and plotted as “naive.” Additionally, we solved the equations with decreasing values of k_1,_ starting with a high value, and referred to this as “pre-activated.” As shown in Fig. 2B, for a range of k_1_<0.85, we found that the Wnt signal strength needed to stabilize β-catenin from the naive state was higher than the signal strength needed to maintain β-catenin once the system had been pre-activated. In contrast, when dephosphorylation of GSK3 by Axin^P^ was omitted from the model, the effect of k1 was identical for “naïve” and “pre-activated” states (Fig. 2C).

Our modeling suggests the β-catenin destruction complex has two stable states, “off” and “on”. This bistability may result from self-perpetuating states such as positive feedback or double-negative feedback (*15*). A bistable system is characterized by two alternative steady states, an off-state and an on-state, without intermediate states (Fig. 3A) (*15*). In a population of cells, the inflection point of the switching will be set by subtle variation in the concentrations and rates of pathway components. This is why, in a uniform sheet of cells, one often observes a salt-and-pepper phenotype rather than perfect collective switching. To simulate the heterogeneous response of a population of cells, we ran the simulation for 2,000 cells where we randomly selected a k_1_ value for each cell from a normal distribution (Fig. S3). We then plotted the distribution of β-catenin with an increase in the mean value of k_1_ (Fig. 3B) that demonstrates the expectation for β-catenin response in a noisy tissue culture system.

**Fig. 3.**
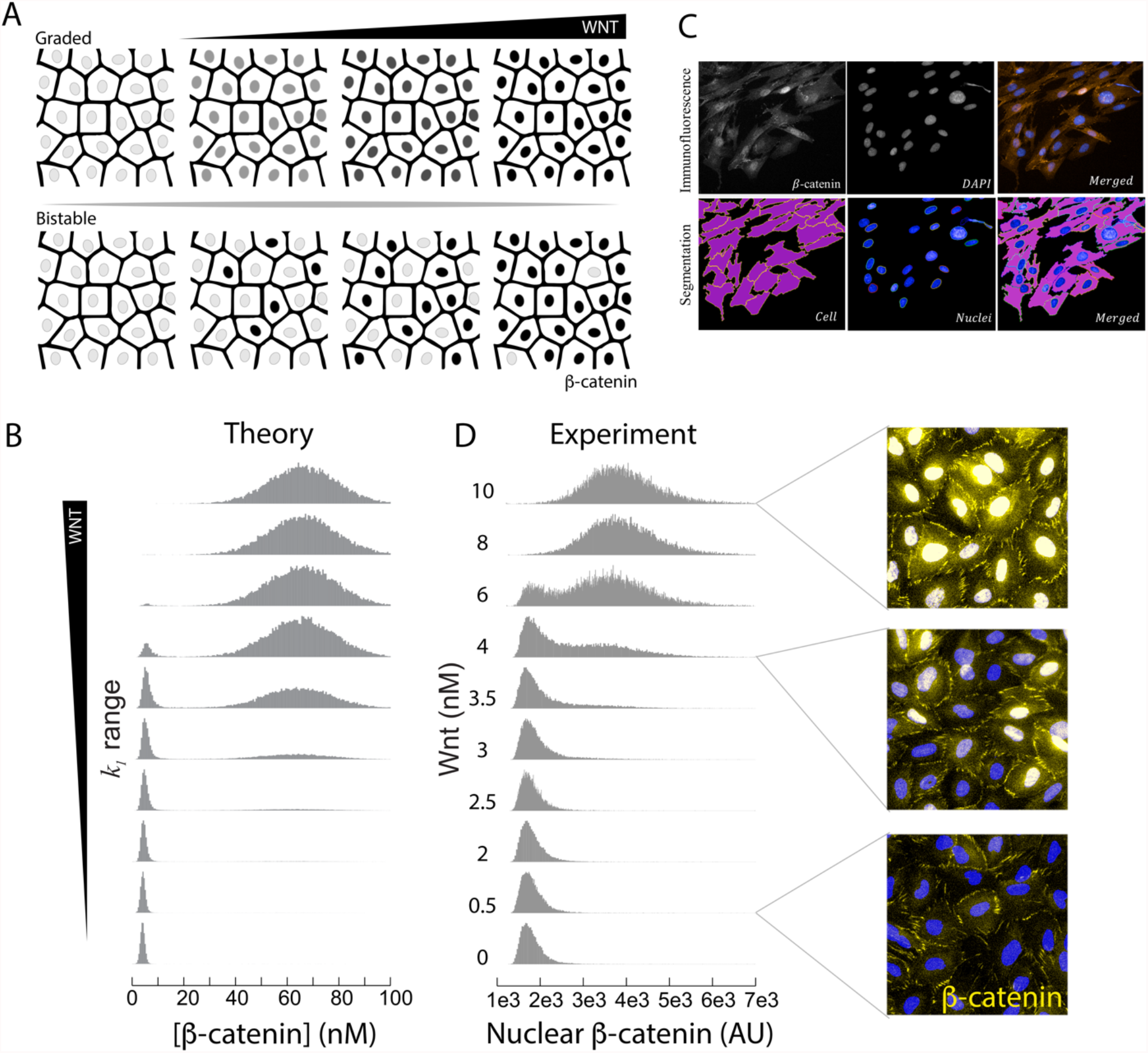
Human colonic epithelial cells respond to Wnt in a bistable manner. (**A**) Depiction of the difference between a graded versus a bistable response in an epithelial monolayer. (**B**,**D**) Density plots of nuclear β-catenin against Wnt concentrations in HCECs under simulated and experimental conditions show a bistable range of 3-6 nM Wnt3a. Steps of automated image processing to quantify nuclear β-catenin immunofluorescence signal. Inserts show representative images of HCECs treated with Wnt at 6 hrs steady state. **(C)** Steps of automated image processing to quantify nuclear β-catenin immunofluorescence signal.

We then validated whether the positive feedback between Axin and GSK3 observed in *Xenopus* egg extracts can lead to a bistable response in mammalian cells activated by Wnt ligands. Accurate single-cell quantification of soluble β-catenin has been challenging due to the high concentrations of non-signaling β-catenin at adherens junctions. We combined automated imaging with custom cell-identification software (Fig. 3C) to analyze primary, immortalized human colonic epithelial cells (HCECs) (*16*). By varying concentrations of purified, recombinant Wnt3a, and measuring nuclear β-catenin, we found that at low Wnt3a concentrations, signaling is in the off-state (nuclear β-catenin is absent), whereas, at high Wnt concentrations, signaling is in the on-state (nuclear β-catenin is present) (Fig. 3D). At intermediate Wnt3a doses, we found a mixed population of cells that were either in the off- or the on-state, with a bistable range between 2 and 6 nM Wnt3a. The β-catenin response to increasing Wnt3a exhibited a Hill-coefficient of ∼6, suggesting cooperativity among responding factors (Fig. S4).

For a system to be genuinely bistable, it must exhibit hysteresis, i.e., the concentration of Wnt needed to induce a given response (after Wnt exposure) is lower than the concentration of Wnt required to mediate the initial response (*15*). An alternative possibility for the salt-and-pepper β-catenin phenotype we observed is the existence of a monostable transcritical bifurcation, a mechanism often found in phase separation settings, in which a two-state system switches states at a single inflection point (i.e., turning on and turning off occur at the same ligand concentration) (*17*). To test this, we fully activated cells with high concentrations of Wnt ligand and then measured the concentrations of Wnt needed to maintain the on-state (Fig. 4A). We observed that cells that were previously treated with a high concentration of Wnt maintained nuclear β-catenin throughout the experiment (Fig. 4B,C) and that this hysteretic behavior persisted several hours after the initial Wnt3a treatment (Fig. S5, S6A).

**Fig. 4.**
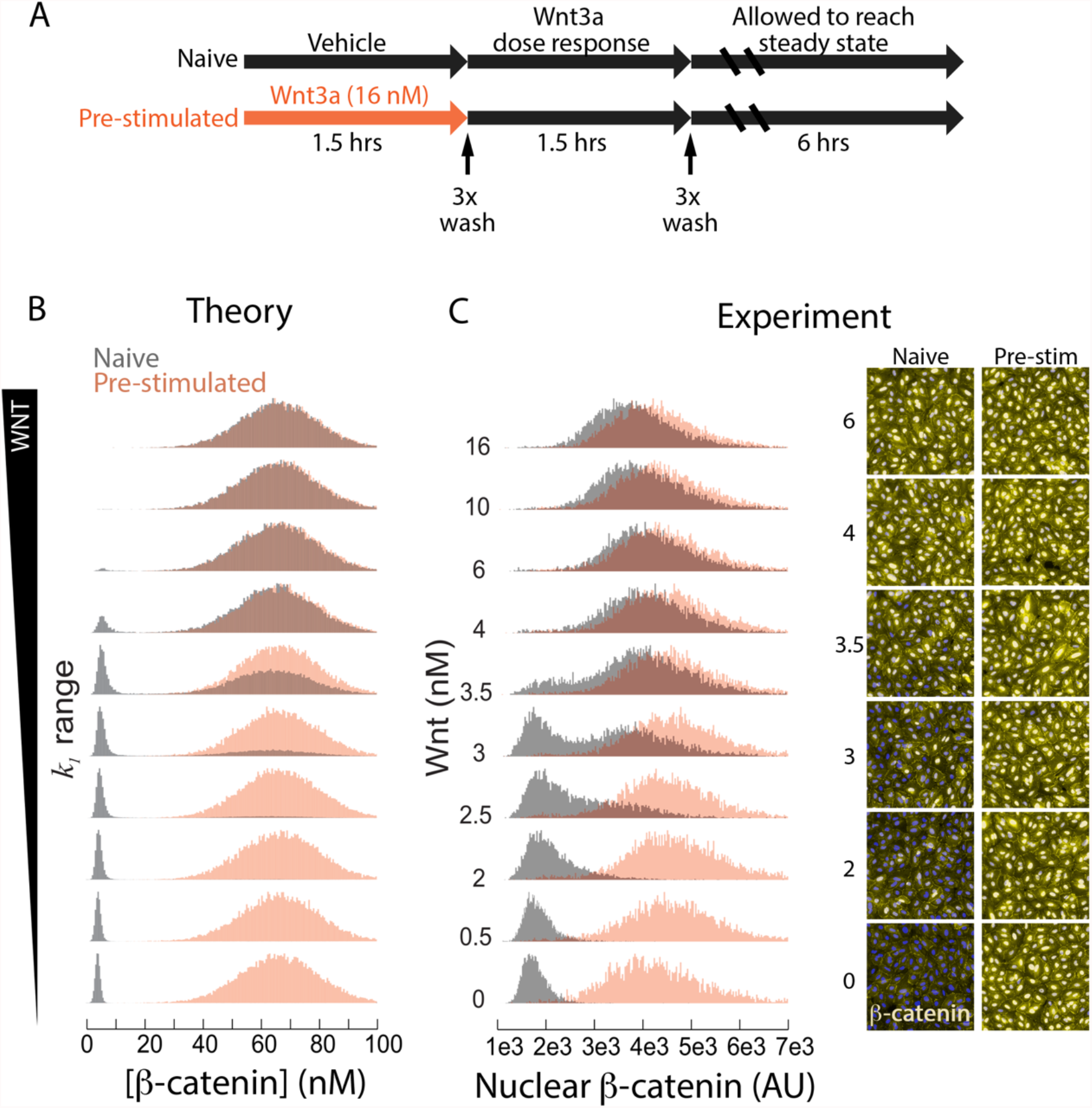
Cells exhibit memory of Wnt stimulation. (**A**) Scheme of the experimental approach to pre-stimulate colonic cells. (**B**,**C**) Model prediction of hysteresis and experimental results from Wnt3a dose-response analyses. Wnt3a dose-response density plots of nuclear β-catenin for HCECs treated with Wnt3a for the first time (naive) or previously pulsed with a high dose of Wnt3a (pre-stimulated). Results from simulated and experimental conditions are shown.

We next tested whether the bistable response observed in HCEC cells in response to Wnt3a treatment was due to positive feedback between Axin and GSK3 revealed in our biochemical experiments (Fig. 1). Our modeling suggested bistability was lost by removing the function of Axin^p^ in the dephosphorylation of GSK3 (Fig. 5A,B). Wnt ligand-mediated activation of the Wnt pathway occurs via a mechanism involving inhibition of GSK3-mediated β-catenin phosphorylation (*14, 18, 19*). We predict, however, that direct inhibition of GSK3 with a small molecule inhibitor that targets its ATP catalytic pocket would break the biochemical GSK3/Axin feedback loop by being insensitive to Axin^p^-dependent dephosphorylation of pS9 GSK3 (Fig. 5C). We treated HCEC cells with the GSK3 inhibitor CHIR99021 (GSK3i) (*20*) and found that, in contrast with the Wnt3a treatment regimen, activation of the pathway with GSK3i treatment failed to promote the bistable behavior of nuclear β-catenin (Fig. 5D). Unlike Wnt3a, the effects of GSK3i on β-catenin nuclear accumulation were readily reversible, and the nuclear β-catenin signal was lost rapidly (without any observable evidence of hysteresis) after the removal of GSK3i (Fig. S6B). Hence, for all GSK3i experiments, we used time points for which we observed a steady-state response after GSK3i treatment, i.e., a minimum of 6 hours treatment (Fig. S7A), but we could not wash out the inhibitor because β-catenin levels rapidly reset to baseline (Fig. S7B). Finally, we observed that GSK3i caused cells to respond in a monostable, graded manner (Fig. 5D, S8). These experimental findings further support the conclusion from our biochemical reconstitution and mathematical model: bistability and hysteresis in Wnt signaling are driven by positive feedback between Axin and GSK3.

**Fig. 5.**
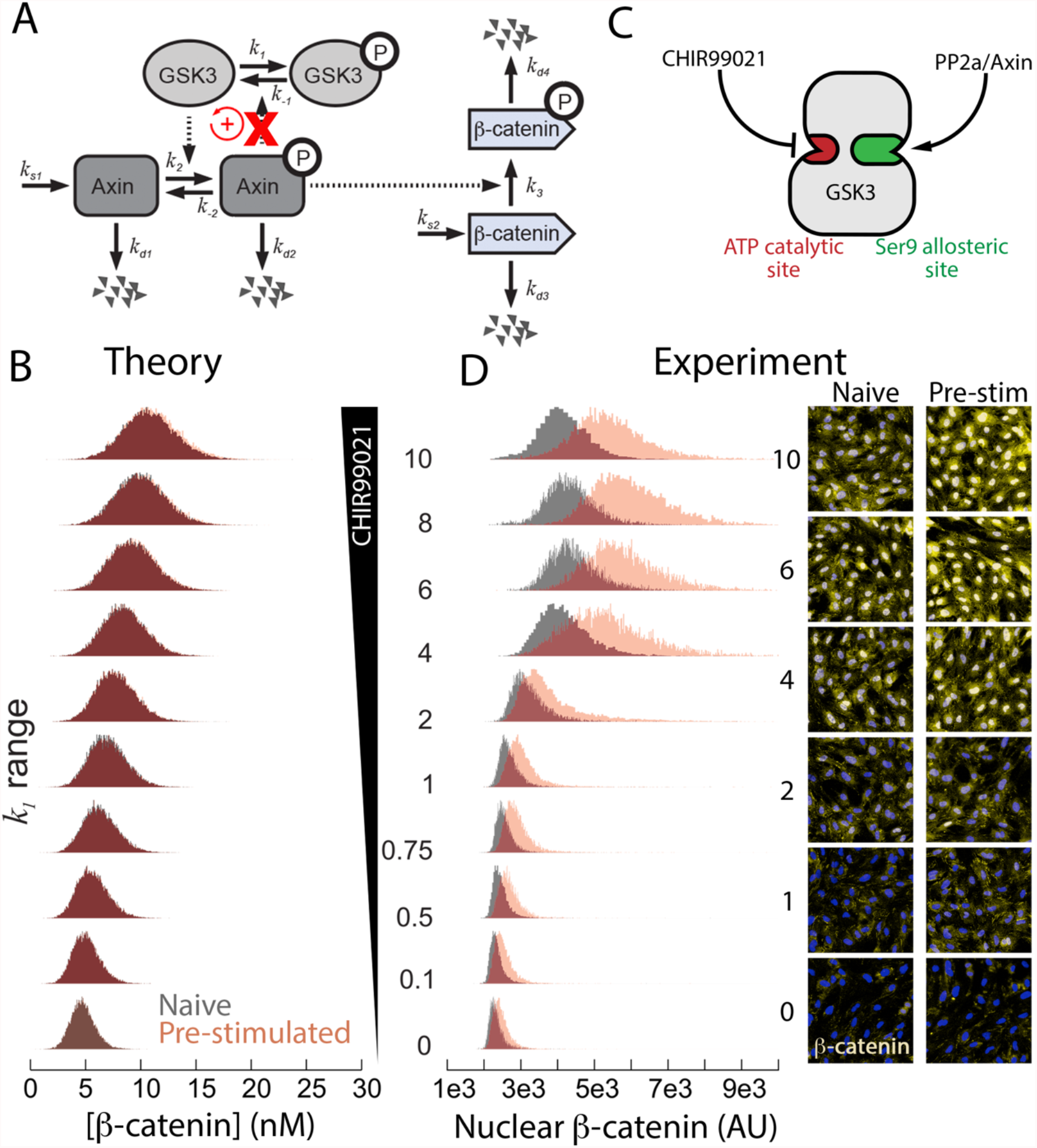
Disrupting positive feedback removes bistability. (**A**) Wiring diagram of β-catenin destruction complex feedback. GSK3i disrupts the positive feedback loop by removing the dependency of Axin^p^ concentration on dephosphorylation of GSK3 (denoted by red X in the diagram) (**B**) CHIR00921 is epistatic to the Axin/PP2a regulation due to direct interaction with the ATP catalytic site on GSK3. (**C**) Model prediction of graded response and experimental results from CHIR009921 dose-response analyses. (**D**) GSK3 inhibition with CHIR99021 treatment results in a graded, monostable response in HCECs.

## DISCUSSION

These experiments demonstrate that a biochemical feedback loop between GSK3 and Axin maintains the β-catenin destruction complex in a stable off- or on-state. This switch-like behavior requires the mutual regulation of GSK3 and Axin via antagonistic behaviors of an additional kinase and phosphatase. We also provided evidence that the PP2A phosphatase acts on GSK3 to remove the inhibitory phosphorylation on serine 9. Modeling of these biochemical events predicted that cells would respond in a binary manner to Wnt pathway stimulation, which was supported by our experiments in human colonic epithelial cells. Additionally, these cells displayed memory to Wnt stimulation, and β-catenin remained in the nucleus even after Wnt ligands had been removed. These results suggest the β-catenin destruction complex displays robustness by existing in two self-sustaining attractor states of active and inactive, which provides a mechanism for suppressing potentially deleterious fluctuations in concentrations and activities of pathway components.

Bistability has emerged as a foundational principle in signal transduction (*21*), yet its existence has been elusive in the Wnt pathway. Beyond suppressing noise within the Wnt pathway, positive feedback in the β-catenin destruction complex provides a mechanism to insulate a pool of GSK3 required in the complex from the total cellular GSK3, thereby preventing crosstalk with other GSK3-regulating pathways such as PI3K/AKT and MAPK (*22, 23*). Furthermore, the existence of both bistable and graded responses could explain why long-range Wnt morphogen activity is dispensable in certain in vivo contexts but essential in others(*24*). The phenomena described herein shed light on a foundational structure of the Wnt/β-catenin pathway that instills robustness and, when perturbed, could lead to vulnerabilities in the accurate processing of Wnt signals.

## Acknowledgments

We are grateful to Dr. Jerry Shay of U.T. Southwestern Medical Center for providing us with the HCEC line. Funding: M.J.C. was supported by the Cancer Biology Training Grant CA T32009213-40. E.A. was supported by The Sidney Hopkins, Mayola B. Vail, and Patricia Ann Hanson Postdoctoral Fellowship. This work was supported by NIH grants GM122516 (E.L.) and CA224188 (E.L. and Y.A.), GM136233 (Y.A.), GM119455 (A.N.K.), DK103126 (C.A.T.), GM134207 (K.D.), and Robert A. Welch Foundation I-1950-20180324. (K.D.)

## Author contributions

Conceptualization, C.A.T., E.L., and K.D. Investigation, M.J.C., E.A., J.R., J.J.T., N.B, A.N.K., Y.A., K.D., E.L., C.A.T.; Writing first draft, M.J.C., C.A.T., E.A., E.L. K.D.; All authors edited the manuscript and approved final submission; Supervision, C.A.T., E.L., and K.D.; Funding acquisition, C.A.T., E.L., and K.D.

## Supplementary Materials for

### Materials and Methods

#### Plasmids and radiolabeled proteins

Radiolabeled β-catenin and Axin were generated in rabbit reticulocyte lysates (Promega) according to the manufacturer’s instructions. Degradation assays were performed based on previously published methods (*7, 25*).

#### *Xenopus* extract studies

*Xenopus* embryos were *in vitro* fertilized, dejellied and extracts prepared as previously described (*7*).

#### Immunoblots

Cells were lysed in non-denaturing buffer (50 mM Tris-Cl, pH 7.4, 300 mM NaCl, 5 mM EDTA, 1% w/v Triton X-100) and the soluble fraction was used for immunoblotting. For Axin immunoblots, Axin was immunoprecipitated with mouse anti-Axin antibody (Zymed) and immunoblotted with anti-Axin 1 goat antibody (R & D). Total GSK3 and GSK3 pS21 (Cell Signaling) were detected from lysates denatured in lysis buffer containing 1% SDS, protease and phosphatase inhibitors.

#### Kinase assays

*In vitro* kinase assays were performed as previously described (*14*).

#### Cell Culture

HEK293 were purchased from ATCC and cultured based in ATCC protocols. Human colonic epithelial cells (HCECs) were cultured in 5% CO_2_ in DMEM supplemented with 10% FBS, 1x penicillin-streptomycin and 1x glutamax.

#### Bistability experiments

HCECs were plated at 20,000 cells/well in fluorescent 96-well plates (Greiner Bio-One; Cat#655090) on day 0. Cells were incubated and allowed to reach 100% confluence. On day 2, cells were treated with increasing concentrations of recombinant human Wnt3A (R&D; Cat#5036-WN-500, with carrier) for 1.5 h. Cells were then washed with PBS thrice and complete media (DMEM high glucose containing 10% FBS, 1x glutamax and 1x penicillin-streptomycin) was added. Cells were incubated for 3 h and fixed with 4% paraformaldehyde-sucrose solution.

##### Hysteresis

HCECs were plated at 20,000 cells/well in fluorescent 96-well plates (Greiner Bio-One; Cat#655090) on day 0 and allowed to reach 100% confluence on day 2. HCECs were treated with Wnt3a long enough to stimulate the pathway (1.5 h), but short enough to avoid negative feedback from the destruction complex (≤ 6 h) (Fig. S6) by Axin2, a transcriptional target of the Wnt pathway. Cells were treated either without (Naive) or with 16 nM of Wnt3A (Pre-stimulated) for 1.5 h. Cells were washed with PBS three times and subsequently treated with increasing concentrations of Wnt3A for 1.5 h. Cells were washed with PBS three times, replaced with complete media for 3 h and fixed.

##### Stimulation through direct GSK3 inhibition (Fig 5)

HCECs were plated as described above. In Fig. S7, cells were treated with CHIR99021 at the indicated concentrations and durations. We chose the concentration of 10 μM for 6h because with these conditions the β-catenin response reached steady state. Cells were treated with increasing concentrations of CHIR99021 (Selleck Chemicals; Cat#S1263) for 6 hrs and then fixed. For the hysteresis experiment using CHIR99021: HCECs were plated as above. On day 2, cells were treated with either DMSO (Naive) or 10 μM of CHIR99021 (Pre-stimulated) for 6 hrs. Cells were washed with PBS three times, treated CHIR99021 dose curve for 6 hrs and fixed.

#### Immunofluorescence

Fixed cells were permeabilized with 0.2% Triton X-100, blocked with 2.5% BSA, and stained with β-catenin antibody at 1:300 (BD Biosciences; Cat#610154) diluted in 2.5% BSA. After washing with PBST (0.1% Tween-20), cells were incubated in secondary antibody conjugated to 1:1000 Alexa fluor (Invitrogen; Cat#A1103) diluted in 2.5% BSA for 2 hrs in the dark. Cells were washed in PBST and DAPI was used to stain nuclei. Cells were imaged on Perkin Elmer Operetta System using a 20x air objective.

#### Nuclear image segmentation

To isolate single cell data, nuclear segmentation was performed using the Perkin Elmer Harmony software as previously described (*26*). β-catenin nuclear intensity was normalized to nuclear area of each cell (nuclear β-catenin). Data analysis codes were custom-built using R.

#### Mathematical modeling

To better understand sufficiency requirements for the emergence of bistability in our system, we developed a minimal mathematical model of the Wnt pathway dynamics. Our equations describe the dynamics of concentrations of Axin, GSK3, and β-catenin in both their phosphorylated and unphosphorylated states. Intermolecular interactions accounted for in the model follow directly from Fig. 2A. Specifically, we assume that Axin is phosphorylated at a rate that increases with the concentration of unphosphorylated GSK3, whereas GSK3 is dephosphorylated at a rate that increases with the concentration of phosphorylated Axin. We further assume that β-catenin is phosphorylated at a rate proportional to the concentration of phosphorylated Axin, which leads to its rapid depletion (since the rate of phospho-β-catenin degradation is assumed to be much higher than that of its unphosphorylated form). Additionally, Axin and β-catenin are produced at constant rates. Finally, Axin and β-catenin undergo degradation in both their phosphorylated and unphosphorylated forms.

Dynamical equations that define our model follow straightforwardly from the above assumptions and are given in Table S1. To analyze the model, we first note β-catenin concentration dynamics do not feedback on the dynamics of the other molecules. In this way, the behavior of β-catenin is a readout of the state of the pathway and need not be considered when examining the nature of its possible dynamical states. Additionally, the total amount of GSK3 is constant (see Eq 4) since its degradation is negligible on the timescale of our experiment (*10*). Thus, three dynamical equations (specifying the dynamics of Axin, phospho-Axin, and unphosphorylated GSK3) suffice to specify the behavior of model uniquely (see Table S1, Eq 1-3).

The key feature of the model is a positive feedback loop between GSK3 dephosphorylation and Axin phosphorylation. Specifically, the rate of GSK3 dephosphorylation increases with the concentration of phosphorylated-Axin (*V*_−1_term), whereas the rate of Axin phosphorylation increases with the concentration of (unphosphorylated) Axin (*V*_2_ term). The activity of the pathway regulates the levels of β-catenin through phosphorylation of β-catenin (*V*_3_ term) which is thereby targeted for proteasomal degradation (*V*_*d*4_ term). The other terms account for Axin-independent phosphorylation of GSK3 (*V*_1_ term), GSK3-independent Axin dephosphorylation (*V*_−2_ term), de novo translation of Axin (*V*_*s*1_ term) and β-catenin (*V*_*s*2_ term), and spontaneous “background” degradation of Axin in its phosphorylated and unphosphorylated states (*V*_*d*2_ and *V*_*d*1_ terms respectively) as well as that of β-catenin in its unphosphorylated state (*V*_*d*3_ term).

In the simplest initial version of the model, it was assumed that no cooperativity was present in any reaction step such that the terms *V*_2_ and *V*_−2_ were linear in [*GSK*3] and [*Axin*^*p*^] respectively. Setting all time-derivatives to zero, these equations are readily solved for the steady state concentrations (which we did using symbolic algebra software Maple, Maplesoft) to obtain two steady state solutions. Setting all time-derivatives to zero, these equations are readily solved for the steady state concentrations (which we did using symbolic algebra software Maple, Maplesoft) to obtain two steady state solutions, one of which is stable and the other unstable. This result indicates that the simplest version of the equations cannot exhibit bistability since bistability requires the coexistence of three steady state solutions (two stable and one unstable). That the simple version of the model does not exhibit bistability can be established rigorously by applying chemical reaction network theory (CRNT) [The Chemical Reaction Network Toolbox, Windows Version | Zenodo].

Hence, to explain bistability, we modified the model to account for the possible presence of cooperativity in both Axin phosphorylation by GSK3 as well as in its dephosphorylation. We note these are assumptions; however, our analysis shows that cooperativity in (at least some of the) intermolecular interactions is required to generate bistable dynamics. The equations are still possible to solve analytically, though the solutions are quite lengthy and thus not presented here. Accounting for possible presence of cooperativity does lead to three solution branches, as indicated in the bifurcation diagram in Fig. S2. To explore the stability of the different branches, we performed numerical simulations where our dynamical equations were solved using an explicit Euler forward scheme. We found that in an open set of parameter values, two stable branches can co-exist with an unstable branch thus implying that our minimal model can account for bistability of the pathway (Fig. S2).

In our minimal model, a steady state with zero concentrations of both phospho-Axin and unphosphorylated GSK3 is present for all values of model parameters. Arguably, this feature is non-generic since it is not present when phospho-GSK3 can be dephosphorylated spontaneously (in the absence of phospho-Axin). However, our model is a good approximation of this later situation if the rate of spontaneous GSK3 dephosphorylation is negligible. It may be shown that a small rate of spontaneous GSK3 dephosphorylation will change the appearance of the bifurcation diagram in Fig. S2 to produce a single root locus where two saddle-node bifurcations are connected by an unstable branch. For high rates of spontaneous GSK3 dephosphorylation, bistability is lost and only a single stable branch will remain. These predictions are in principle testable, provided one can control the rate of GSK3 dephosphorylation for example by a phosphatase, presumably PP2a. However, this work is beyond the scope of the present study.

To further examine the dynamics of the pathway and to more closely compare theoretical predictions to experimental results, we examined the response of the pathway to quasi-static variation of a control parameter. Specifically, we varied the rate of spontaneous GSK3 phosphorylation (k_1_) and examined the resulting steady-state concentration of β-catenin. This was done using two procedures that we term “Naive” and “Pre-stimulated”. In the former case, the value of k_1_ was initially set to a low value and was then increased quasi-statically. In the latter case, k_1_ was gradually decreased, starting with a large initial value. Fig. 2 shows the plots of the steady-state β-catenin levels as a function of k_1_ corresponding to the two simulated protocols. It is seen that the two curves do not coincide, indicating the presence of hysteresis. Specifically, in the Pre-stimulated case when k_1_ was large initially, the initial levels of β-catenin were correspondingly high and remained high for a substantial range of (decreasing) k_1_ values. In the Naive case, when k_1_ was increased from a value initially set to be low, steady-state β-catenin levels remained much lower than seen in the Pre-stimulated case (for the same values of k_1_). This hysteretic behavior is a key hallmark of bistability and is readily anticipated from the phase portrait given in Fig. S2.

We next asked if our minimal computational model can interpret our observations on the distribution of Wnt pathway activation levels in cell populations (Fig. 3D, 4C, 5D). Clearly, the precise level of nuclear β-catenin varies from cell to cell. To capture this variation, we considered an ensemble of cells, with each cell having a different random value of spontaneous GSK3 phosphorylation k_1_. Expectedly, this results in a stochastic distribution of steady-state β-catenin levels. Next, we examined changes in the simulated distribution of pathway activation as the mean value of k_1_ was varied according to the two protocols described above and as shown in Fig. 2. We found that the distribution of β-catenin levels becomes bimodal within a finite intermediate range of the mean k_1_ values. This is another key hallmark of bistability and is in complete agreement with the experimental data (see Fig. 4B and 4C). At the same time, average β-catenin levels follow the same qualitative trend as was seen in Fig. S9, where the deterministic dynamics without stochastic variation in k_1_ were examined. Finally, we asked if the dynamics of the pathway may be rendered monostable by means of perturbations that interfere with the positive feedback between Axin and GSK3. To this end, we made the rate of Axin phosphorylation independent of GSK3 concentration. In this case, the dynamics became monostable such that no hysteresis was seen when k_1_ was first increased and then decreased quasi-statically, see Fig. S9E.

To account for the effect of Wnt stimulation on the pathway, we considered four alternative scenarios. Specifically, in our model, Wnt can stabilize β-catenin by 1) increasing *k*_1_, 2) decreasing *k*_−1_, 3) increasing *k*_−2_ or 4) decreasing *k*_2_. Expectedly, in all these cases, the system exhibits hysteresis (Fig. S9), since, generically, the presence of hysteresis does not depend on the particular choice of the control parameter (as long as varying the control parameter allows to move from a bistable to a monostable regime).

**Table S1:**
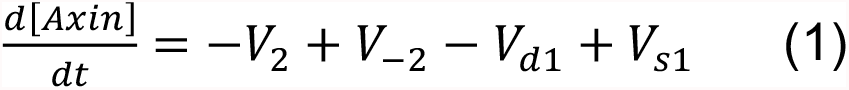
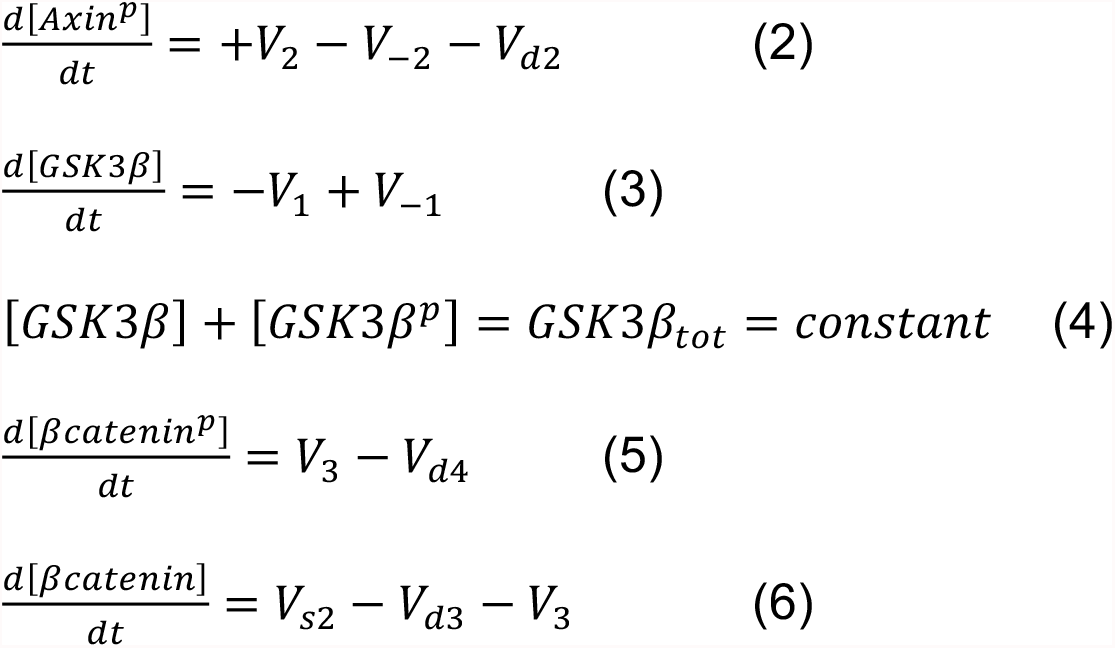
Set of ordinary differential equations for the system in Fig. 2A.

**Table S2:**
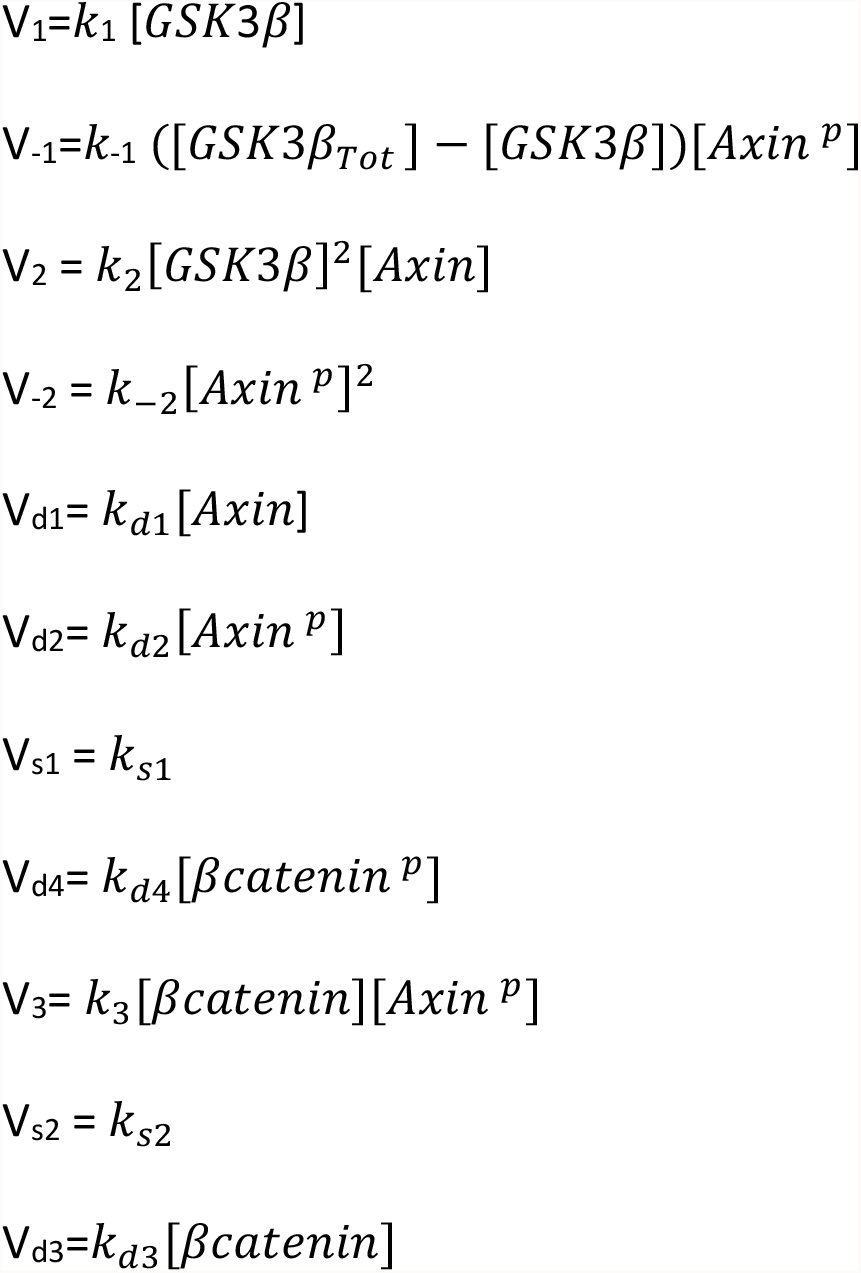
Reaction rates for the model.

**Table S3:** Rate constants used in the model.

| Variable | Value | Definition | Ref |
| --- | --- | --- | --- |
| $k_{s1}$ | $8.2 \times 10^{-5} \text{ nM min}^{-1}$ | Axin synthesis rate | (10) |
| $k_{d1}$ | $0.167 \text{ min}^{-1}$ | Axin degradation rate constant | (10) |
| $k_{d2}$ | $0.004 \text{ min}^{-1}$ | Axin<sup>p</sup> degradation rate constant | Assumption |
| $k_1$ | Fig S9A & Fig 2B : $0.05\text{-}2 \text{ [min}^{-1}]$<br>Fig S9B: $1 \text{ [min}^{-1}]$<br>Fig. S9C: $1 \text{ [min}^{-1}]$<br>Fig. S9D: $1 \text{ [min}^{-1}]$<br>Fig. S9E & Fig 2C: $0.05\text{-}4 \text{ [min}^{-1}]$ | phosphorylation rate of GSK3 | |
| $k_{-1}$ | Fig S9A & Fig 2B : $1 \text{ [nM}^{-1} \text{ min}^{-1}]$<br>Fig S9B: $0.05\text{-}4 \text{ [nM}^{-1} \text{ min}^{-1}]$<br>Fig. S9C: $1 \text{ [nM}^{-1} \text{ min}^{-1}]$<br>Fig. S9D: $2 \text{ [nM}^{-1} \text{ min}^{-1}]$<br>Fig. S9E & Fig 2C: $0.001 \text{ [min}^{-1}]$ | Dephosphorylation rate of GSK3<sup>p</sup> | |
| $k_2$ | Fig S9A & Fig 2B : $2 \text{ [nM}^{-2} \text{ min}^{-1}]$<br>Fig S9B: $2 \text{ [nM}^{-2} \text{ min}^{-1}]$<br>Fig. S9C: $0.05\text{-}10 \text{ [nM}^{-2} \text{ min}^{-1}]$<br>Fig. S9D: $2 \text{ [nM}^{-2} \text{ min}^{-1}]$<br>Fig. S9E & Fig 2C: $2 \text{ [nM}^{-2} \text{ min}^{-1}]$ | phosphorylation rate of Axin | |
| $k_{-2}$ | Fig S9A & Fig 2B : $2 \text{ [nM}^{-1} \text{ min}^{-1}]$<br>Fig S9B: $2 \text{ [nM}^{-1} \text{ min}^{-1}]$<br>Fig. S9C: $2 \text{ [nM}^{-1} \text{ min}^{-1}]$<br>Fig. S9D: $0.05\text{-}10 \text{ [nM}^{-1} \text{ min}^{-1}]$<br>Fig. S9E & Fig 2C: $2 \text{ [nM}^{-1} \text{ min}^{-1}]$ | Dephosphorylation rate of Axin<sup>p</sup> | |
| $k_3$ | $5\text{nM}^{-1} \text{ min}^{-1}$ | Phosphorylation rate of β-catenin | Assumption |
| $k_{s2}$ | $0.42 \text{ nM min}^{-1}$ | Synthesis rate of β-catenin | (10) |
| $k_{d3}$ | $6.3 \times 10^{-3} \text{ min}^{-1}$ | Degradation rate of βcatenin | (10) |
| $k_{d4}$ | $0.42 \text{ min}^{-1}$ | Degradation rate of β-catenin<sup>p</sup> | (10) |
| $GSK3_{Tot}$ | 50nM | Total concentration of GSK3 | (10) |

**Table S4:** Two initial conditions used to solve the ODEs.

| Experiment[nM] | Axin | Axin<sup>p</sup> | GSK3 | βcatenin | βcatenin<sup>p</sup> |
| --- | --- | --- | --- | --- | --- |
| Naive | $4 \times 10^{-4}$ | $5 \times 10^{-3}$ | 50 | 1 | 30 |
| Pre-stimulated | $6.1 \times 10^{-4}$ | $1 \times 10^{-6}$ | 2 | 30 | 1 |

**Fig S1.**
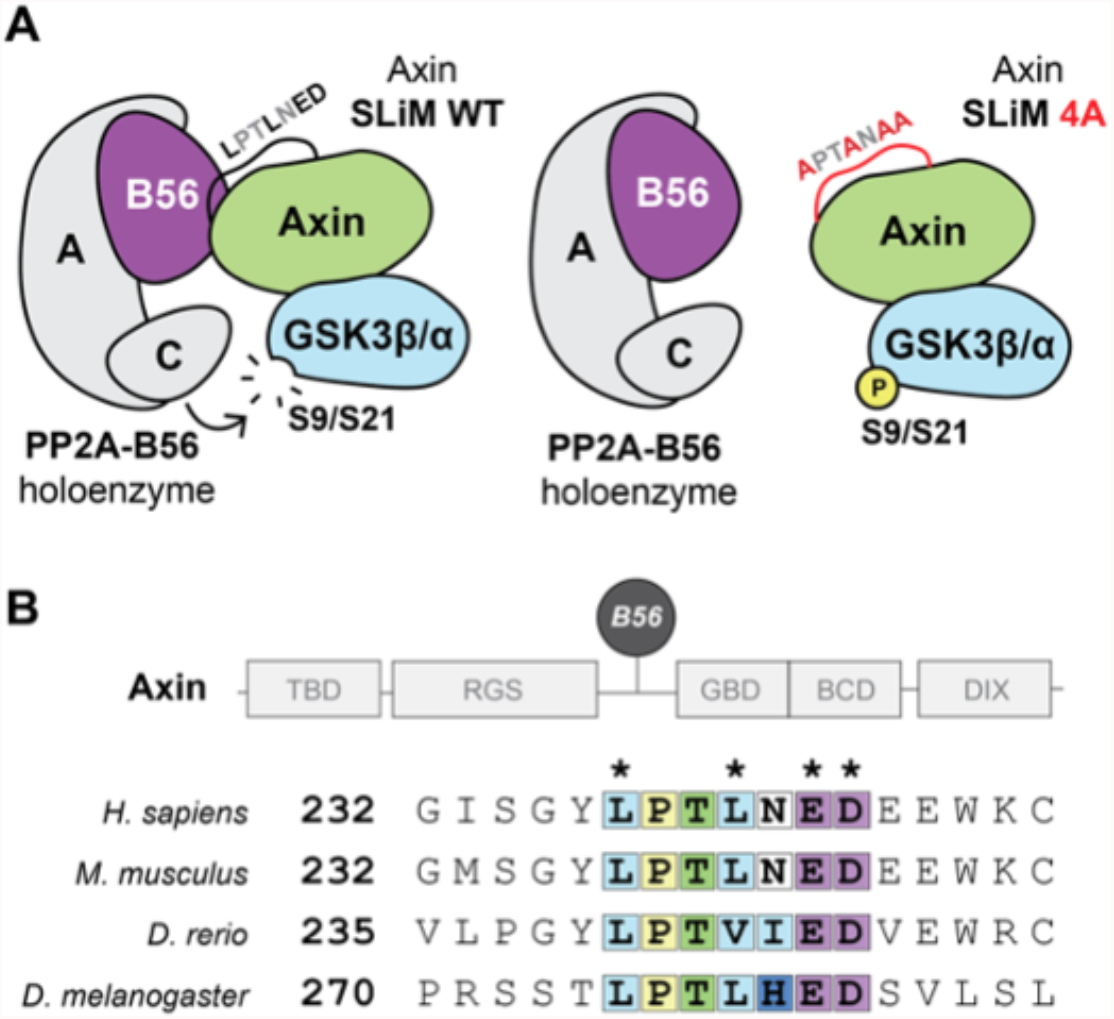
(**A**) Cartoon of Axin SLiM site mutant effect on GSK3 activation. (**B**) Alignment of the B56 binding site sequence in four species reveals strong evolutionary conservation.

**Fig S2.**
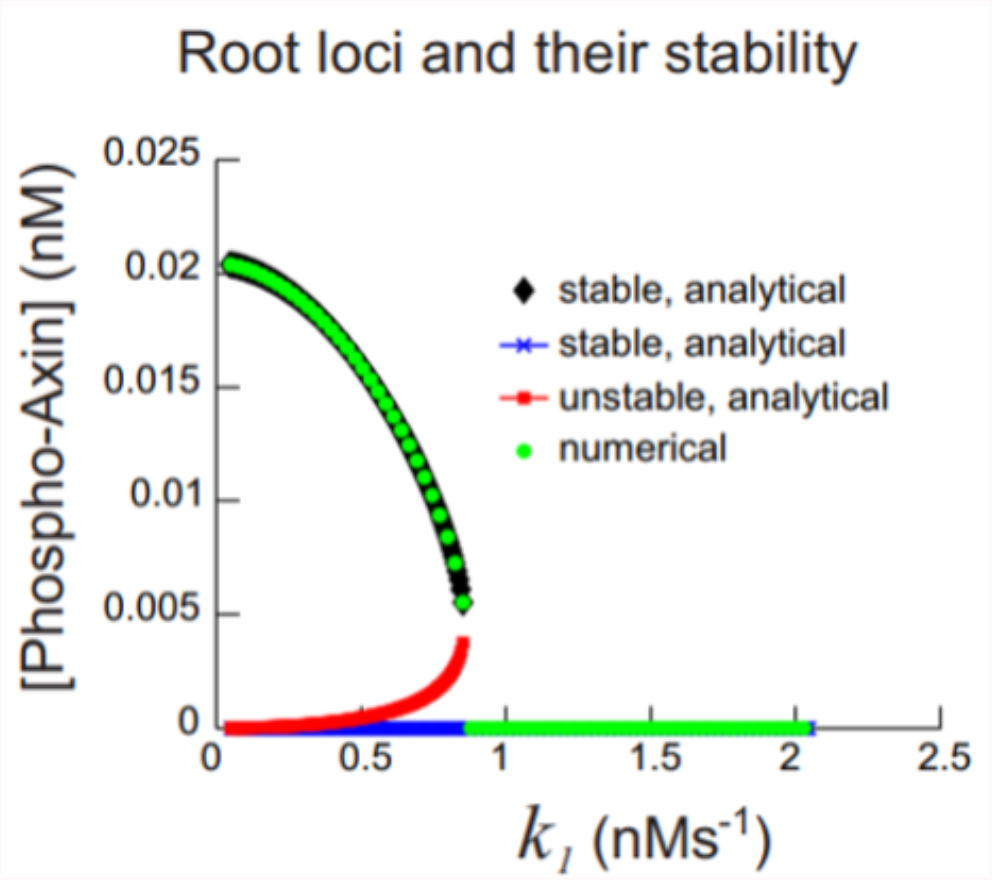
Solving the ODEs analytically leads to three fixed points. We explored the stability of the different branches using numerical simulations where our dynamical equations were solved. Two of the fixed points were stable and can co-exist while the third fixed point was unstable.

**Fig S3.**
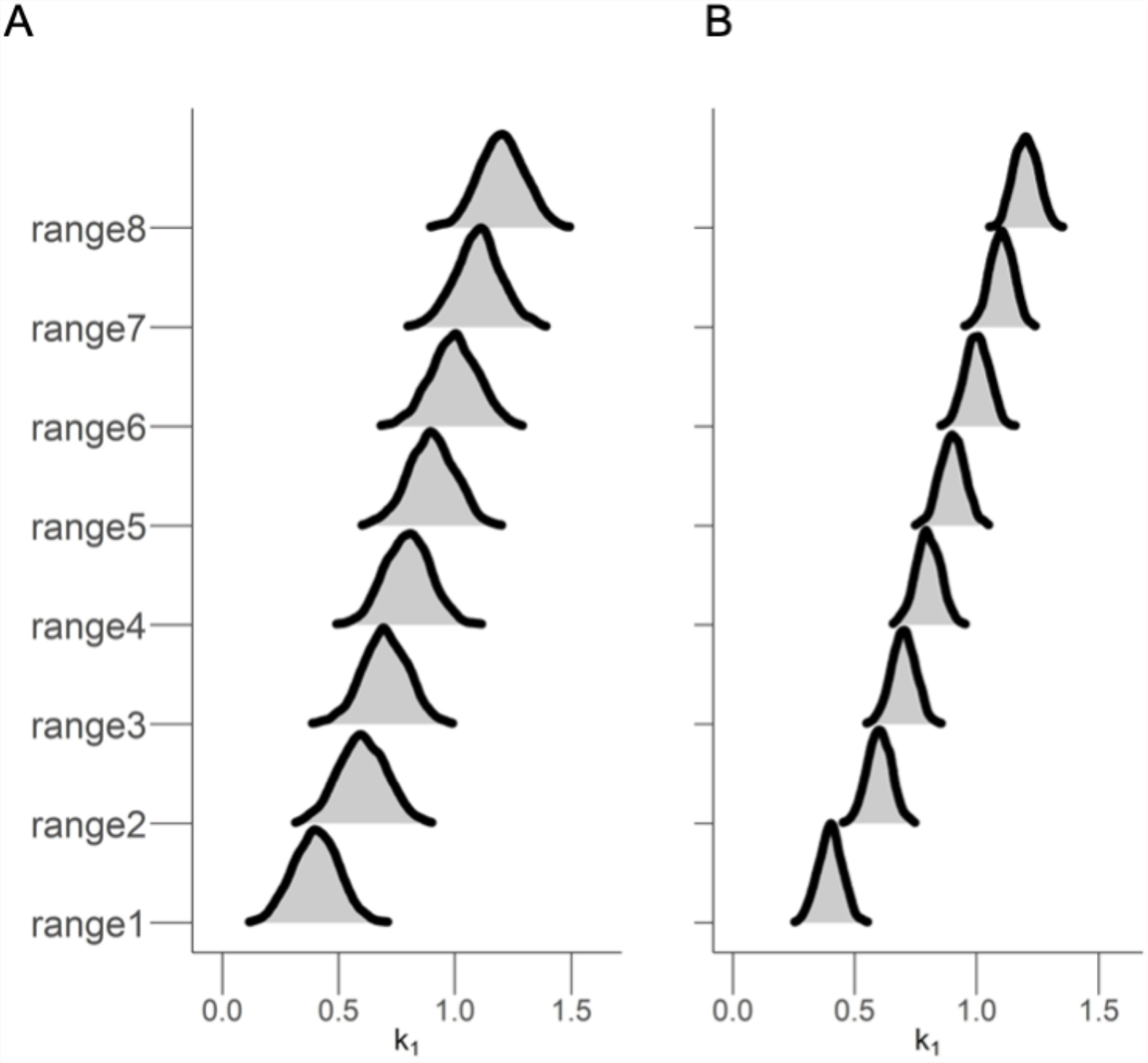
To account for heterogeneity of the cells, we ran the simulation for 2000 cells and in each run we randomly choose a k_1_ value from a normal distribution instead of a single value. (**A**) Range of k_1_ used for the model in which there is a positive feedback loop and bistability response is observed as shown in Fig. 3B and Fig. 4B. (**B**) Range of k_1_ used for the model where the positive feedback loop is broken, and we see a graded response as shown in Fig. 5D.

**Fig S4.**
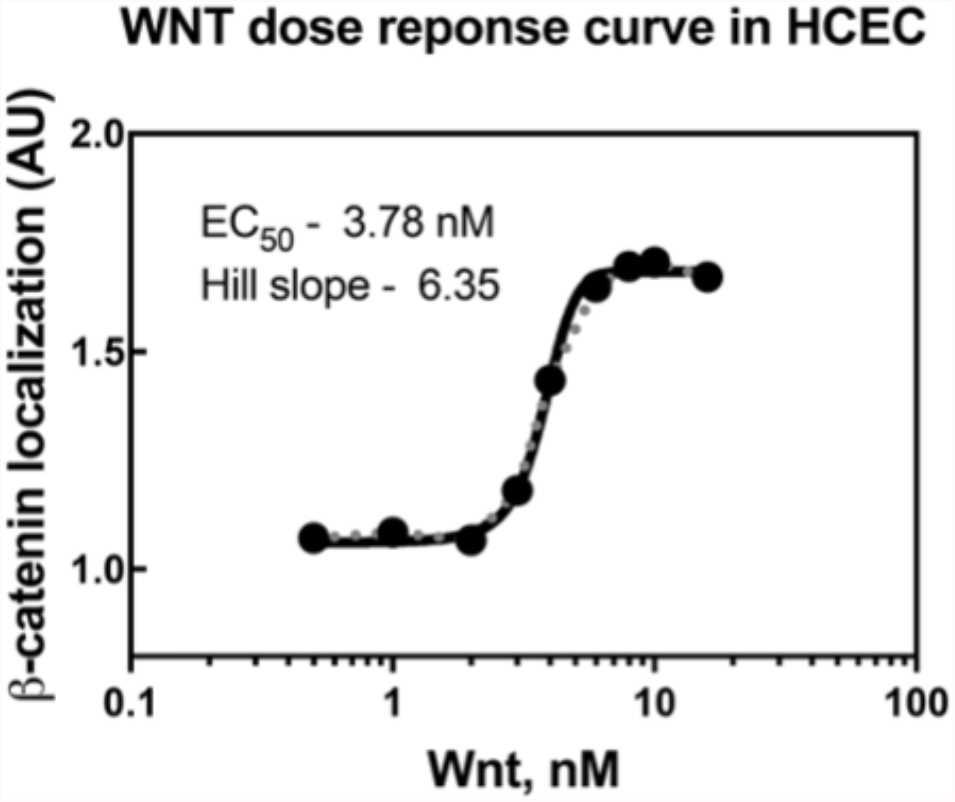
Plot of mean β-catenin response to increasing and decreasing Wnt3a stimulation from Fig. 3D, (EC_50_= 3.78 nM; n_H_ = 6.35).

**Fig S5.**
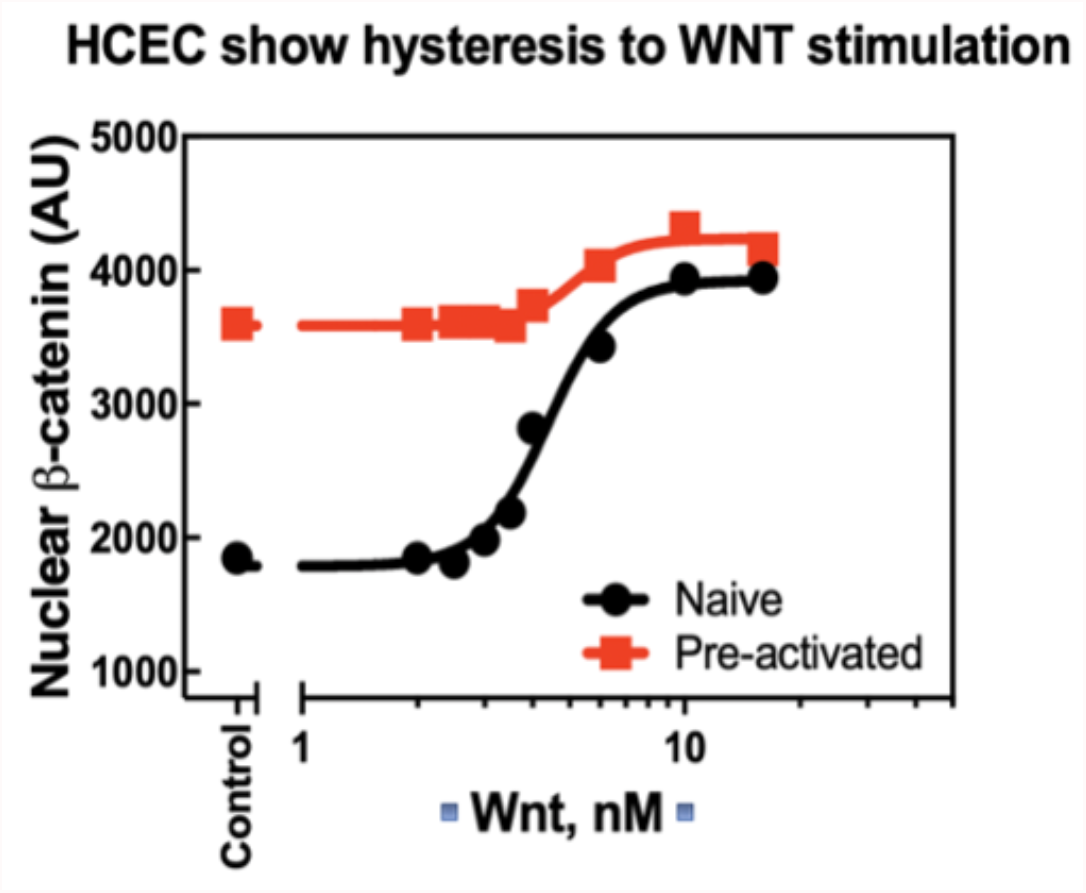
Plot of mean β-catenin response to increasing and decreasing Wnt3a stimulation from Fig. 4D.

**Fig S6.**
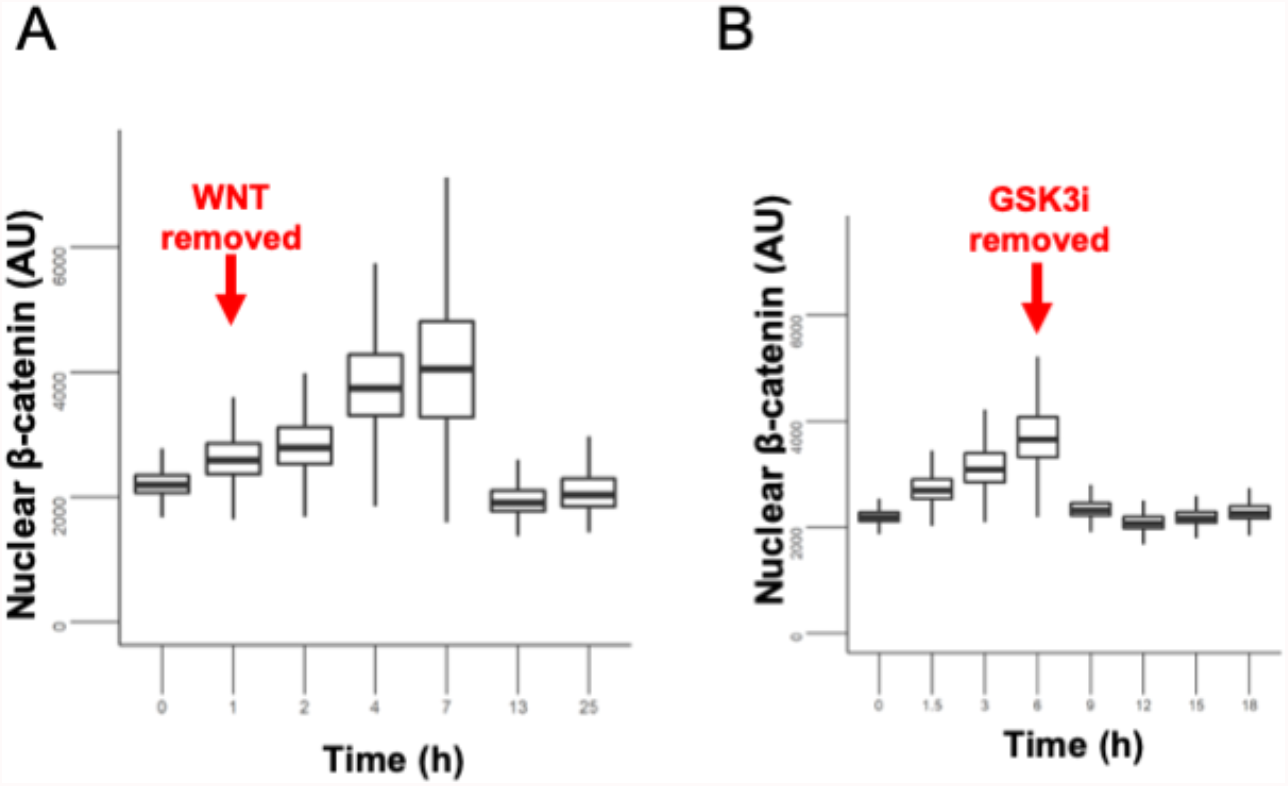
(**A**) HCECs were treated with recombinant Wnt3a (16 nM) for one hours, washed three times and regular growth media minus Wnt3a was added back. Nuclear β-catenin is plotted. Nuclear β-catenin remains high at 6 h after removal but afterward comes back down. (**B**) HCECs were treated with different concentrations of the CHIR99021 for various durations. Cells were allowed to reach steady-state using nuclear localization of β-catenin as a readout. The effect of CHIR99021 on nuclear β-catenin localization in HCECs is reversible 3h after removal of the compound. CHIR99021 was used at 10 μM concentration for six hours.

**Fig S7.**
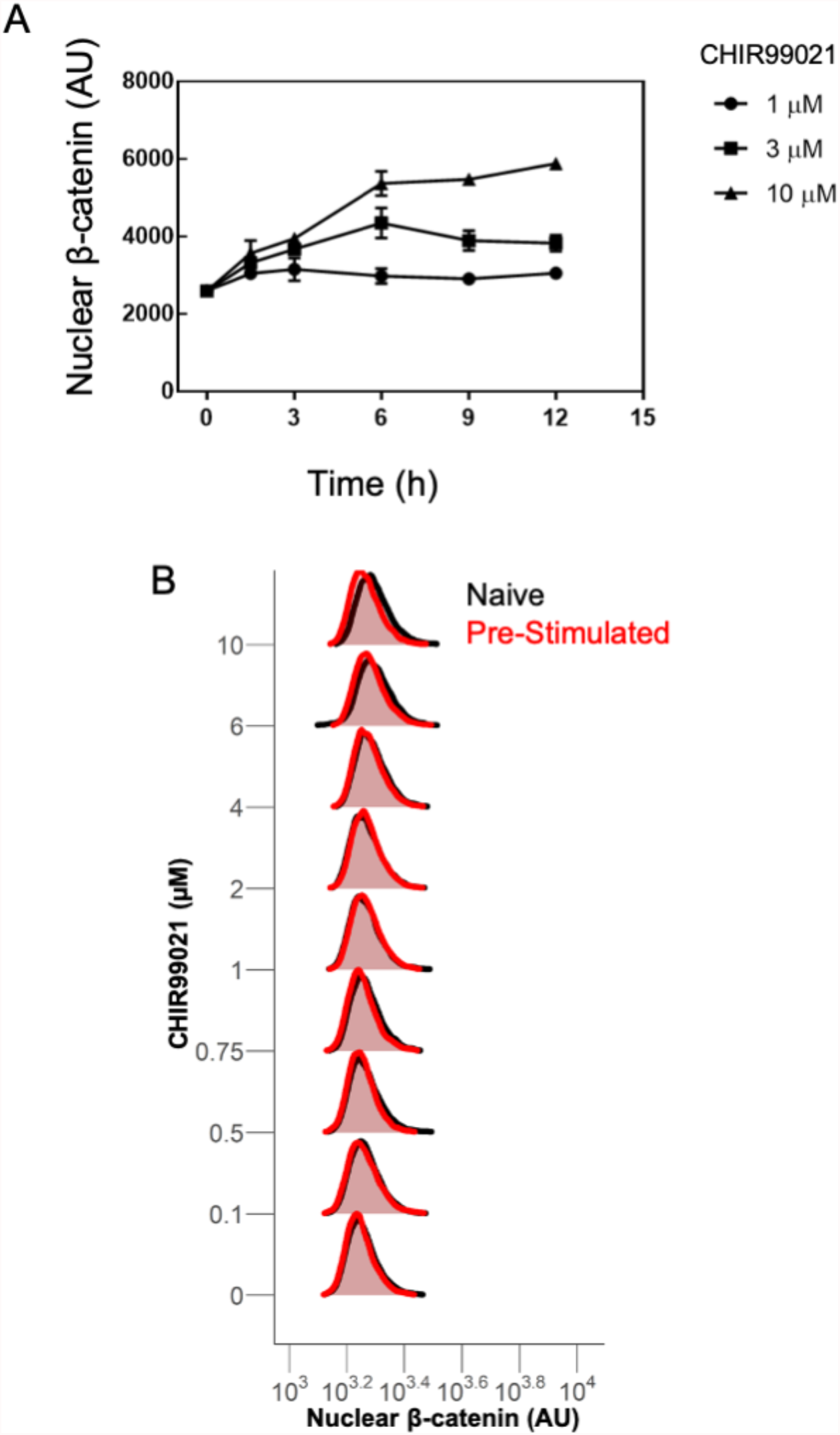
(**A**) HCECs were treated with different concentrations of GSK3i CHIR99021 for various durations. Nuclear localization of β-catenin is plotted. β-catenin steady state was achieved at 6 hours under sustained CHIR99021 stimulation. (**B**) β-catenin hysteresis experiment was performed with identical treatment times as was described in Fig. 4A, except CHIR99021 was used instead of Wnt3a. Unlike the effect observed in cells treated with Wnt3a, cells treated with CHIR99021 cannot maintain memory of the stimulation and return to basal β-catenin concentrations at the 6 hr timepoint.

**Fig S8.**
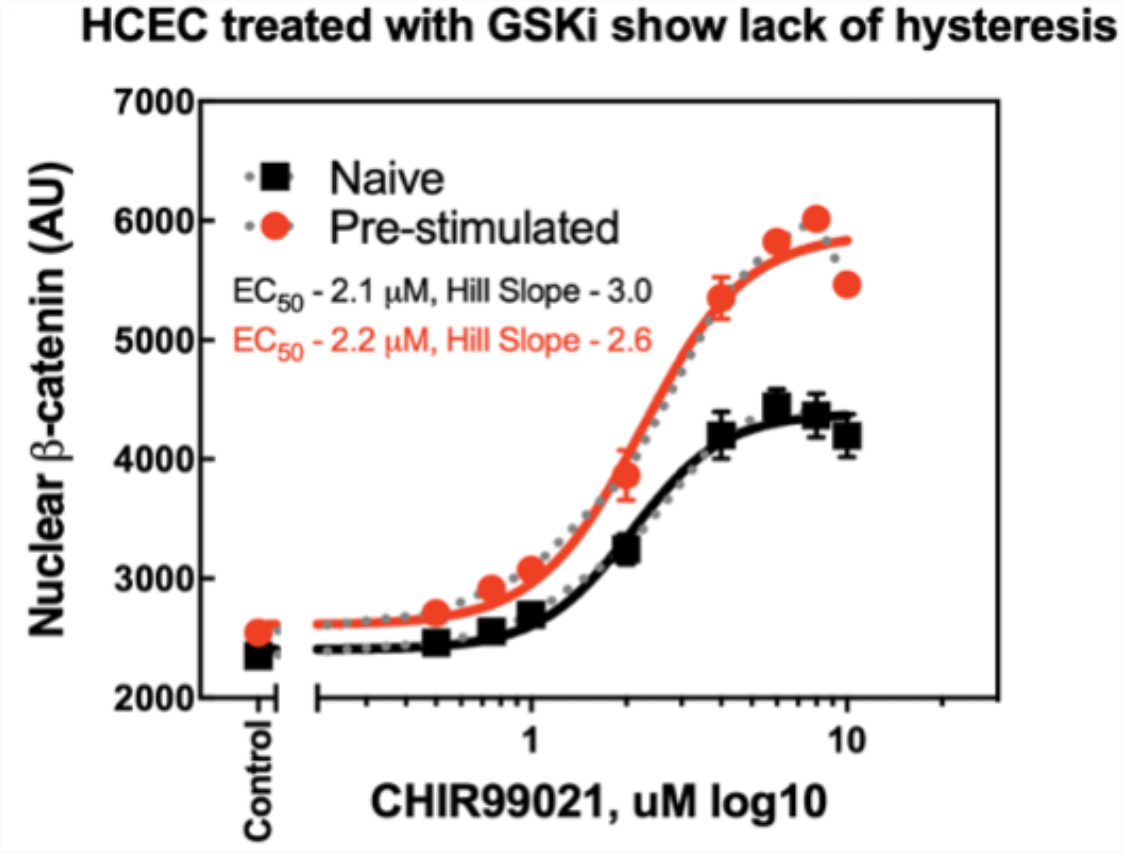
Plot of mean β-catenin response to increasing and decreasing CHIR99021 stimulation from Fig. 5D.

**Fig S9.**
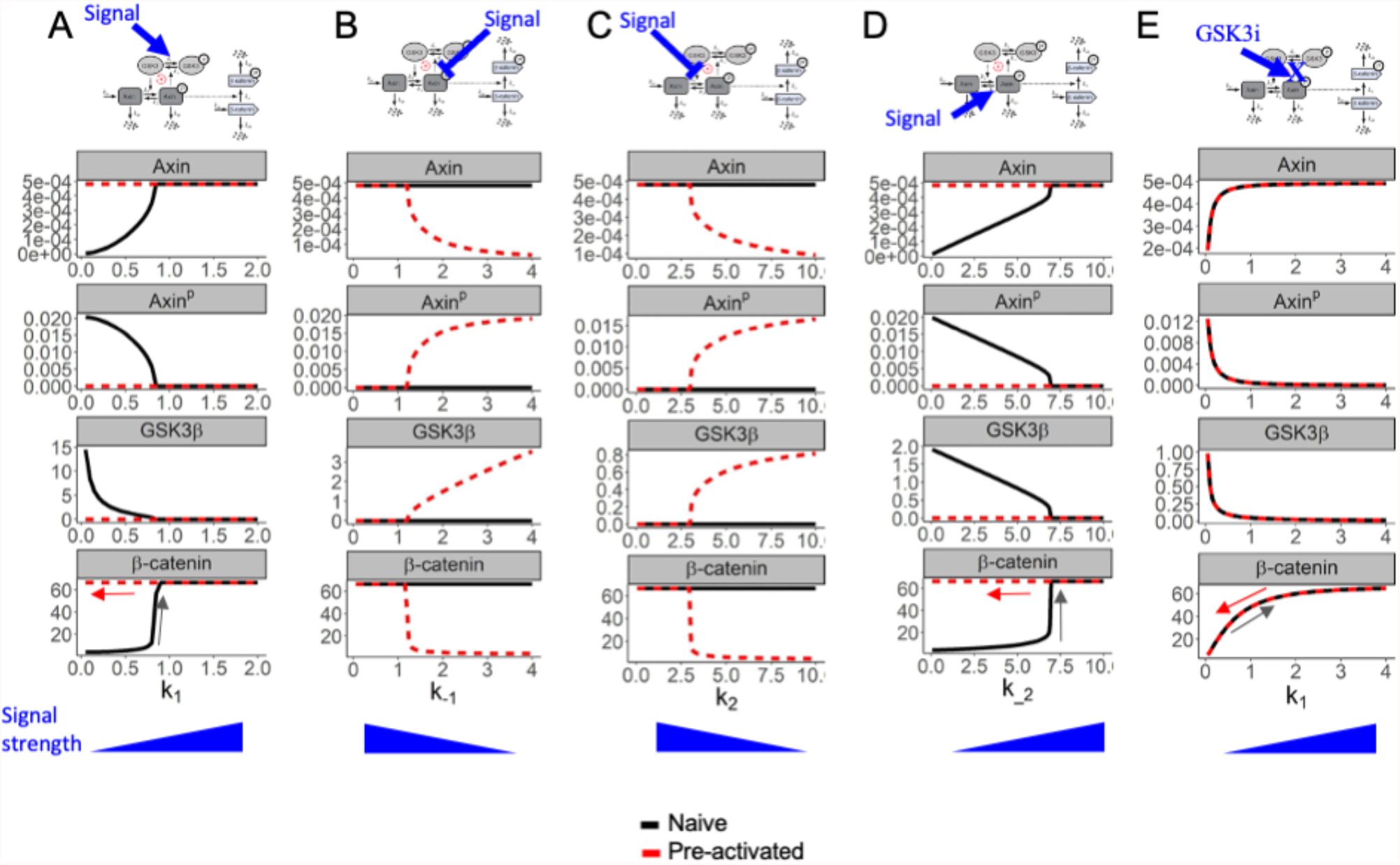
We assumed four different scenarios where in each scenario one of k1, k2, k-2, and k-1 rate constants changes and the others remain constant with increasing Wnt. (**A**) Wnt increases k1 with k_2=2, k2=2, and k_1=1. (**B**) Wnt decreases k-1 with k_2=2, k2=2, and k1=1. (**C**) Wnt decreases k2 with k_2=2, k1=1, k_1=1. (**D**) Finally Wnt increases k-2 with k2=2, k1=1,k_1=2. As shown in these four panels for a range of these parameters we see bistability in the concentrations of the molecules. (**E**) We simulated adding GSK3i as disrupting the positive feedback loop and leading to a graded response in β-catenin concentration rather than a bistable response.

